# The Infectious Bronchitis Virus Coronavirus Envelope Protein Alters Golgi pH to Protect Spike Protein and Promote Release of Infectious Virus

**DOI:** 10.1101/440628

**Authors:** Jason W. Westerbeck, Carolyn E. Machamer

## Abstract

Coronaviruses (CoVs) are important human pathogens with significant zoonotic potential. Progress has been made toward identifying potential vaccine candidates for highly pathogenic human CoVs, including use of attenuated viruses that lack the CoV envelope (E) protein or express E mutants. However, no approved vaccines or anti-viral therapeutics exist. CoVs assemble by budding into the lumen of the early Golgi prior to exocytosis. The small CoV E protein plays roles in assembly, virion release, and pathogenesis. CoV E has a single hydrophobic domain (HD), is targeted to Golgi membranes, and has cation channel activity *in vitro*. The E protein from the avian infectious bronchitis virus (IBV) has dramatic effects on the secretory system, which requires residues in the HD. Mutation of the HD of IBV E during infection results in impaired growth kinetics, impaired release of infectious virions, accumulation of IBV S protein on the plasma membrane when compared IBV WT infected cells, and aberrant cleavage of IBV S on the surface of virions. We previously reported the formation of two distinct oligomeric pools of IBV E in transfected and infected cells. Disruption of the secretory pathway by IBV E correlates with a form that is likely monomeric, suggesting that the effects on the secretory pathway are independent of E ion channel activity. Here, we present evidence suggesting that the monomeric form of IBV E correlates with a rise in the pH of the Golgi lumen. We demonstrate that infection with IBV induces neutralization of Golgi luminal pH, promoting a model in which IBV E alters the secretory pathway through interaction with host cells factors, protecting IBV spike protein (S) from premature cleavage and leading to the efficient release of infectious virus from the cells.

## Introduction

The majority of human coronaviruses (CoVs) cause mild disease phenotypes. However, when novel coronaviruses like severe acute respiratory syndrome (SARS)-CoV and Middle East respiratory syndrome (MERS)-CoV emerge from their animal reservoirs to infect humans they elicit a robust and aberrant immune response that can lead to a very serious and deadly pneumonia (1–3). Importantly, there are no effective vaccines or therapeutics to treat these CoVs. Efforts to develop long-term therapeutic strategies to combat novel, highly pathogenic CoVs will be aided by increased understanding of conserved viral mechanisms at the level of their cell and molecular biology.

One of the more fascinating and relatively enigmatic aspects of CoV biology is that CoV virions bud into the lumen of the secretory pathway at the endoplasmic reticulum-Golgi intermediate compartment (ERGIC), and then must navigate through the Golgi and the anterograde endomembrane system to be efficiently released from the host cell. It has been demonstrated that the structure and function of the Golgi depends upon an acidic pH gradient that decreases from the lumen of the *cis*-Golgi to the lumen of the *trans*-Golgi. This pH gradient is produced by a balance maintained by proton influx into the lumen of the Golgi, proton leak, and counter-ion conductance (4). Pharmacological manipulations of the pH gradient, resulting in neutralization of the lumen, have all been shown to cause slow trafficking of cargo through the Golgi as well as alteration in Golgi morphology (4–10). A class of small viral membrane proteins with ion channel activity, called viroporins, have been shown to have dramatic effects on the secretory pathway, similar to those elicited by pharmacological manipulation of luminal pH (11–21). Several well-studied members of this viroporin family of proteins include the Influenza A M2 protein (IAV M2), hepatitis C virus (HCV) p7 protein, and the CoV envelope (E) protein. These representative viroporins demonstrate several common functional features despite differences in viral assembly and budding locations. It has been suggested the role of M2 in the secretory pathway is to neutralize luminal pH to protect the HA fusion protein of influenza from premature activation (5–7). Overexpression of M2 causes secretory pathway disruption where the rate of intracellular trafficking is slowed and Golgi morphology is altered (8). HCV p7 is also thought to play a protective role by allowing egress of viral structural proteins through the secretory pathway. HCV can be partially rescued by both pharmacological neutralization of the luminal spaces by bafilomycin A1 and by *in trans* expression of IAV M2 (9, 10). Similar to M2, the infectious bronchitis virus (IBV) coronavirus E protein elicits multiple secretory pathway disruption phenotypes when overexpressed in mammalian cells.

To understand CoV E at a cell biological level, a recombinant virus system was used to replace the hydrophobic domain (HD) of IBV E with the HD of the vesicular stomatitis virus glycoprotein (VSV G), and the recombinant virus was called IBV EG3 (22, 23). Replacing the HD of IBV E with a heterologous sequence of the same length does not impair Golgi targeting or interaction with IBV M during assembly (24, 25), but would be expected to impair ion channel function. One-step growth curves revealed that IBV EG3 virus grew to a titer 10-fold lower than IBV WT virus in infected Vero cells. At late times post-infection, the majority of infectious virus resides in the supernatant surrounding IBV WT infected cells, while the majority of infectious IBV EG3 virus is intracellular (23). Vero cells infected with IBV EG3 accumulate more IBV S protein on the plasma membrane than IBV WT infected cells and this accumulation of IBV S leads to an increase in the size and rate of formation of the virus-induced syncytia (23). Most virions purified from IBV EG3 infected cells lack a full complement of spikes and S is aberrantly cleaved near the virion envelope, potentially explaining the reduced infectivity of released particles (22). A build-up of vacuole-like compartments containing virions as well as other aberrant material in IBV EG3 infected cells may explain the damage to S (22, 23).

Intriguingly, when IBV WT E is transiently overexpressed in HeLa cells, the Golgi completely disassembles while the Golgi in cells overexpressing IBV EG3 is intact (23). This observation suggested that IBV E alters the secretory pathway of the host cell. Expression of IBV E or EG3 reduces trafficking rates of both membrane and secretory cargo (23). Given that release of infectious IBV EG3 is reduced, it was surprising that wild-type E protein reduced cargo trafficking. It was hypothesized that since the HD was required, alteration of the lumen by E ion channel activity was required for maintaining intact virus, and the reduced rates of trafficking were an acceptable compromise for the virus.

Studies probing the nature of CoV E ion channel activity have centered on understanding the residues required for this activity and the associated pathogenic and cell biological phenotypes elicited by different CoV E proteins. Two residues in the HD of SARS-CoV E, N15 and V25, have been shown to promote viral fitness during infection (11, 21). Mutation of N15 or V25 abolishes ion-channel activity of SARS-CoV E in artificial membranes (11, 21). We previously reported that the E protein of IBV is expressed in mammalian cells is found in two pools by velocity gradient analysis: a low molecular weight pool (LMW) and a high molecular weight pool (HMW) (26). The LMW pool represents IBV E in a monomeric state while the HMW pool correlates with a homo-oligomer of IBV E. When mutations corresponding to the conserved HD residues of SARS-CoV E that inhibit ion channel activity were made in IBV E (T16A and A26F), we found that the HD mutants segregated primarily into one oligomeric pool or the other. The E^T16A^ mutant was primarily in the HMW pool while the E^A26F^ mutant was found primarily in the LMW pool. The presence of the LMW pool of IBV E, the predominant and likely monomeric form found when E^A26F^ is present, correlates with the secretory pathway disruption associated with the WT IBV E protein (26). This was surprising in that it suggested an E ion channel-independent role for IBV E associated with manipulation of the secretory pathway. It was recently reported that that these HD mutants do abolish ion channel activity of IBV E in artificial membranes, and virus titers are reduced by a log in the supernatant of infected cells, suggesting a defect in virion release (27). Our data on the IBV-EG3 virus corroborates this study (23).

Herein, we provide evidence for the neutralization of the Golgi during IBV infection and we demonstrate that transient over-expression of the IBV E protein, but not HD mutants deficient in the LMW pool of IBV E, is sufficient to cause a significant increase in the pH of the Golgi lumen. We suggest that increased trafficking and aberrant cleavage of the IBV S protein observed during IBV EG3 infection may reflect the detrimental effect of normal pH on IBV S processing. We demonstrate that IBV S processing and trafficking is similarly aberrant when coexpressed with EG3 but not WT E, and that IAV M2 can substitute for WT E to protect S from premature cleavage.

## Results

### IBV induces an increase in Golgi luminal pH during infection

Given that the effects of IBV E overexpression on the Golgi are similar to those in cells where the luminal pH is neutralized, we first measured luminal pH in IBV-infected cells. We used flow cytometric analysis of a ratiometric pHluorin molecule targeted to the Golgi with a reporter consisting of the green fluorescent protein (GFP) pHluorin molecule fused to the membrane targeting sequence of the TGN38 *trans*-Golgi network resident protein (28). We chose to use the *trans*-Golgi network pHluorin because the TGN is the most acidic compartment of the Golgi and thus any alteration in pH would likely be most detectable in this compartment. We generated a clonal Vero cell line that stably expressed the TGN38-pHluorin (Figure 1A). The generation of this cell line allowed us to ensure that all infected cells were expressing the TGN38-pHluorin. The cells were treated with cycloheximide for 60 minutes to chase newly synthesized TGN38-pHluorin from the ER. To generate a pH calibration curve, cells expressing the Golgi pHluorin molecule were subjected to treatment with buffers ranging from pH 5.5 to 7.5 in the presence of the ionophores monensin and nigericin prior to flow analysis (Figure 1B). The emission ratios of the biphasic pH-sensitive pHluorin at these known pH values can then be used to predict the Golgi luminal pH in cells infected with IBV in buffer at physiological pH and lacking ionophores (Figure 1C). Infection resulted in a robust increase in the Golgi luminal pH.

**Figure 1.**
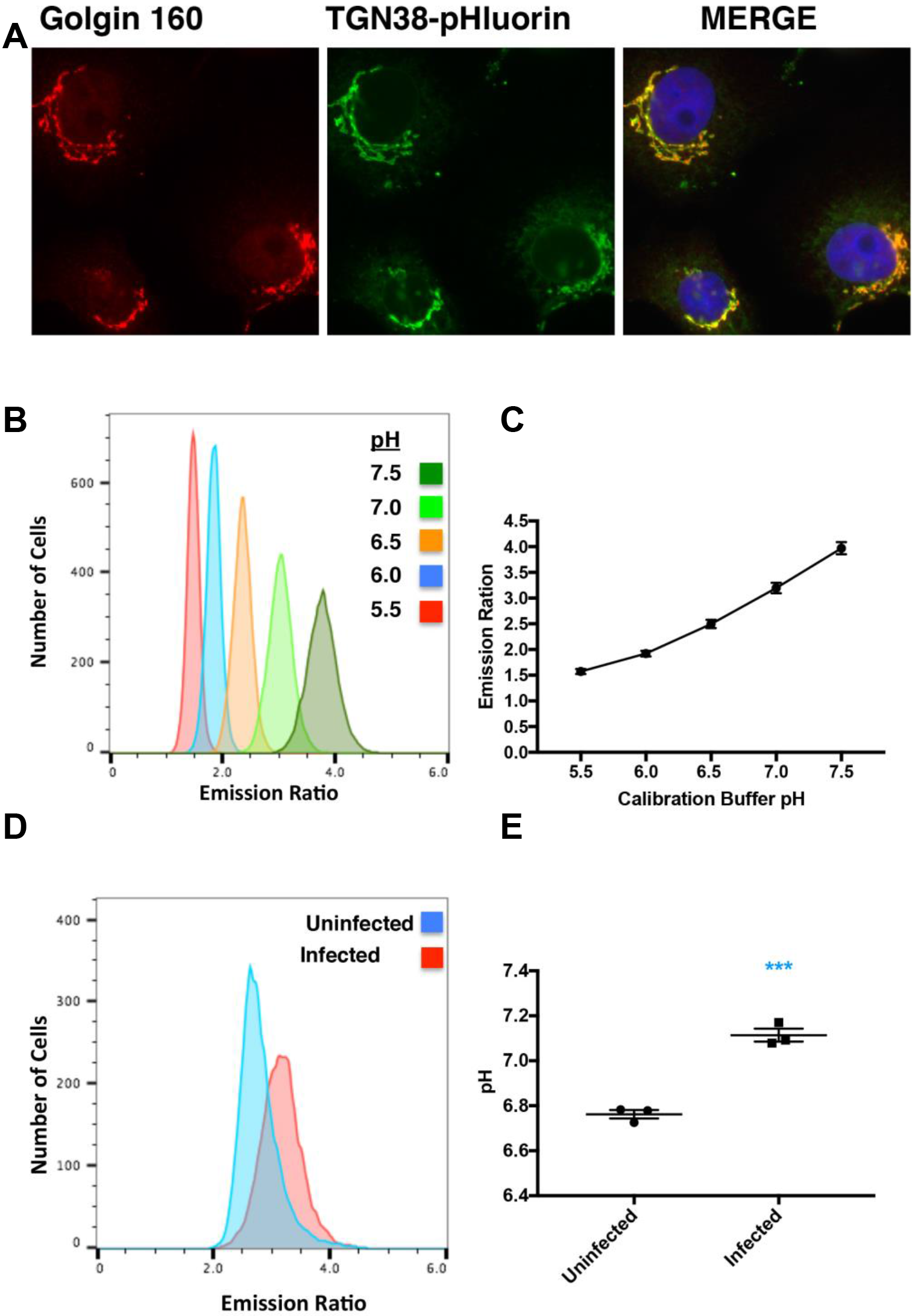
IBV infection alters Golgi pH. (A) Vero cells stably expressing TGN38-pHluorin were evaluated by indirect immunofluorescence microscopy. Cells were labeled with rabbit anti-golgin160 and mouse anti-GFP. (B) The pH of Vero cells stably expressing the TGN38-pHluorin was assessed by determining the ratio of the pH sensitive dual emission spectrum by flow cytometry. The cells were in buffers of known pH and contained ionophores to equilibrate the extracellular and luminal pH of the Golgi. A representative flow cytometry experiment is graphed. (C) Calibration curves were generated from data like that illustrated in (B), in order to calculate the pH of cells infected with IBV or uninfected cells. The calibration curve pictured was derived from n=3 independent experiments (~10,000 cells each). Error bars = SEM. (D) The cell emission ratios for Vero cells infected or mock infected with IBV and stably expressing TGN38-pHluorin from a representative experiment are pictured. (E) The average calculated pH values from n=3 independent experiments (~10,000 cells each) are graphed. Unpaired t-tests were performed in Prism at 99% confidence, with an assumption of equal variance. Two or more stars were considered to represent a significant difference. ***P≤0.001. Error bars= SEM.

**Figure 2.**
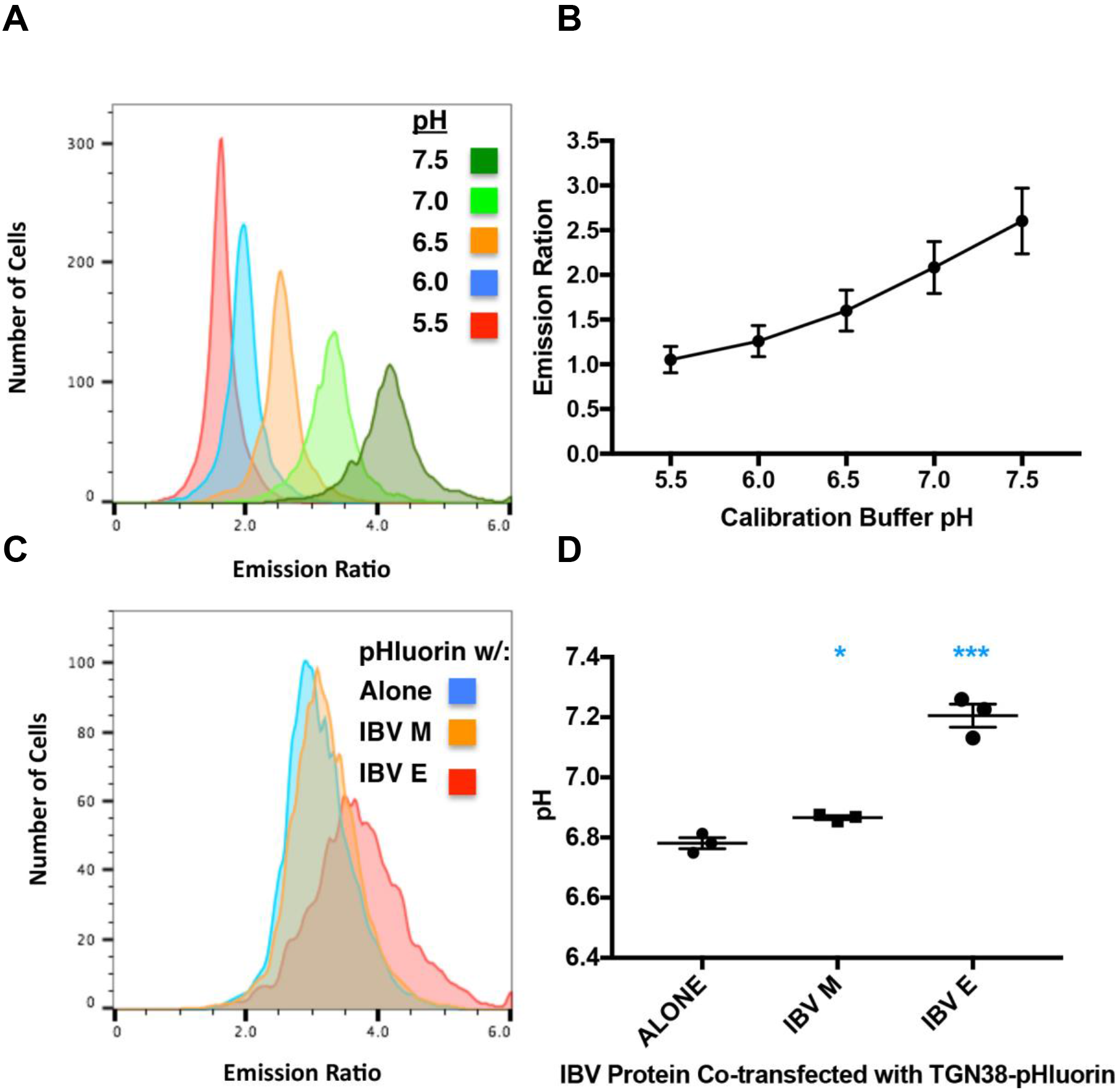
Over expression of IBV E alters Golgi pH.(A) HeLa cells transiently expressing the TGN38-pHluorin were assessed by determining the ratio of the pH sensitive dual emission spectrum by flow cytometry. The cells were in buffers of known pH and contained ionophores to equilibrate the extracellular and luminal pH of the Golgi. A representative flow cytometry experiment is graphed. (B) Calibration curves were generated from data like that illustrated in (A), in order to calculate the pH of cells expressing IBV E or E mutants, in combination with the TGN38-phluorin. The calibration curve pictured was derived from n=7 independent experiments (~5,000 cells each). Error bars = SEM. (C) The cell emission ratio for HeLa cells expressing TGN38-pHluorin alone and with IBV E or IBV M (control) from a representative experiment are pictured. (D) The average calculated pH values from n=3 independent experiments are graphed (~5,000 cells each). Unpaired t-tests were performed in Prism at 99% confidence, with an assumption of equal variance. Two or more stars were considered to represent a significant difference. * P≤0.05; ** P≤0.01; ***P≤0.001. Error bars= SEM.

We attempted to measure the Golgi luminal pH in cells infected with IBV-EG3, but were unable to achieve a high percent of infected cells in the absence of syncytium formation, since this virus is not efficiently released and spreads best by cell-cell fusion. We found that syncytia were fragile, and this precluded flow cytometric analysis. Instead we turned to transfected cells to determine if the E protein could neutralize the Golgi lumen.

### Overexpression of the IBV E protein increases the pH of the Golgi lumen

To determine if the pH change in infected cells was caused by the E protein, HeLa cells were cotransfected with a plasmid encoding IBV E along with the pHlorin-TGN38 reporter. We used transient transfection of the reporter here to ensure the pHluorin expressing cells were also expressing the E protein. In separate cells, we included vector alone and a plasmid encoding IBV M (as another overexpressed Golgi membrane protein) as controls. As described above, transfected cells were pretreated with cycloheximde for 60 min to chase newly synthesized pHlorin-TGN38 out of the ER. A standard curve was produced in cells treated with ionophores in calibrated pH buffers. As shown in Figs 2B and C, IBV E neutralized the trans-Golgi luminal pH when over expressed in HeLa cells, whereas the IBV M protein did not.

### The increase in Golgi pH correlates with the LMW pool of IBV E

To determine the role of IBV E oligomerization in the alteration of Golgi luminal pH, we analyzed two HD point mutants of IBV E that segregate into different oligomeric states. Our previous findings suggest that IBV E^A26F^ is found predominantly in the LMW, likely monomeric form, and IBV E^T16A^ is found predominantly in the HMW, higher-order oligomer (26). In addition, we analyzed the EG3 mutant of IBV E, in which the entire HD of IBV E was replaced by a heterologous HD sequence. Both IBV E^T16A^ and IBV EG3 had pH measurements similar to the IBV M membrane protein control (pH 6.95 and pH 6.87, respectively), while IBV E^A26F^ elicited a robust pH increase similar to that of the wild-type IBV E protein (pH 7.18, Figure 3A.) This suggested that the LMW pool of IBV E correlates with the luminal pH increase of the Golgi in addition to the secretory pathway disruption demonstrated in our previous work (23, 26, 29). The same experiment was performed with a *medial-Golgi* pHluorin, GnT1-pHluorin (Gnt1, N-acetylglucosaminyltransferase I), to assess if the alteration in pH was specific to the TGN. Indeed, both IBV E and IBV E^A26F^ elicited a robust pH increase (Figure 3B). Interesting, IBV E^T16A^ increased the pH significantly, though not as robustly as IBV E and IBV E^A26F^. We found that the pH of the trans-Golgi (measured with TGN38-pHluorin) was higher than the *medial-Golgi* (measured with GnT1-pHluorin). This was unexpected and is addressed in the discussion. The results in transfected cells implicate the monomeric form of IBV E in neutralization of the Golgi lumen during infection and transfection.

**Figure 3.**
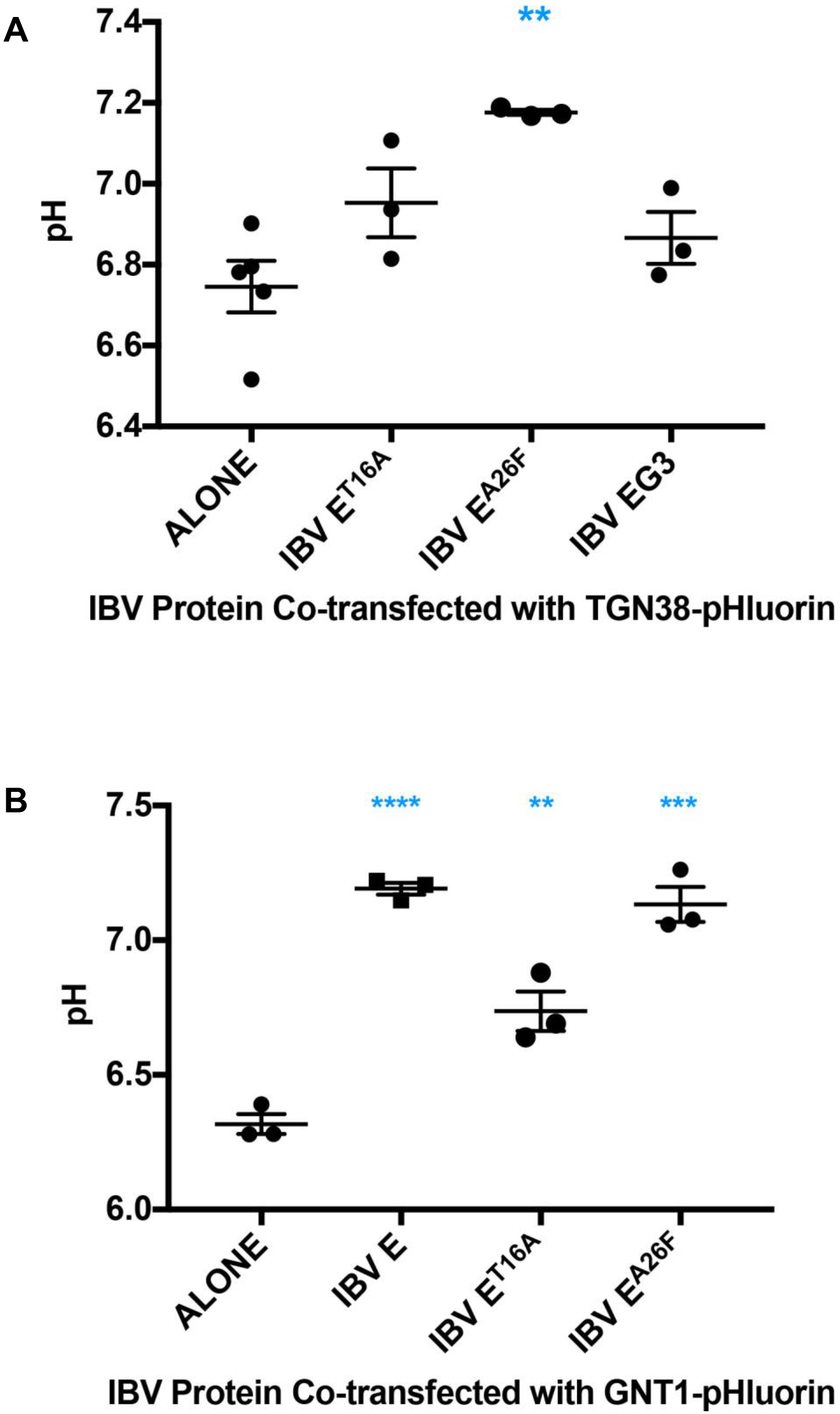
Change in Golgi pH correlates with LMW HD mutant of IBV E. (A) HeLa cells transiently expressing the TGN38-pHluorin with IBV E or HD mutants were evaluated by flow cytometry and the average calculated pH values from n=3 independent experiments are graphed (~5,000 cells each). (B) HeLa cells transiently expressing a *medial-Golgi* tagged pHluorin, GnT 1-pHluorin, with IBV E or HD mutants were evaluated by flow cytometry and the average calculated pH values from n=3 independent experiments are graphed (~5,000 cells each). Unpaired t-tests were performed in Prism at 99% confidence, with an assumption of equal variance. Two or more stars were considered to represent a significant difference. ** P≤0.01; ***P≤0.001. Error bars= SEM.

### Attempts to rescue IBV EG3 by manipulation of Golgi pH

Despite intense efforts we were unable to conclusively determine whether an increase in Golgi pH could rescue the deficiencies of the IBV EG3 virus. We tried drugs that neutralize acidic compartments (baflinomycin A1, monensin and ammonium chloride), as well as overexpression of influenza A M2, a pH activated proton channel (4). However, the drugs inhibited exocytosis at all concentrations used (during short or long infections), and, we were unable to obtain a high percentage of M2-transfected cells that were subsequently infected with IBV-EG3. In several experiments where the percent of transfected and infected cells was greater than 40%, we obtained 40-75% increases in release of infectious IBV-EG3 virus, but most experiments failed to show a reasonable overlap of transfection and infection and rescue of IBV-EG3 infectivity (data not shown). Additionally, attempts to make stable lines expressing M2 and pHluorin did not yield lines expressing M2 at a high enough level to alter the Golgi pH. We thus turned to another approach to assess the role of neutralization of the Golgi by IBV E during infection.

### Expression of IAV M2 decreases the total amount cleaved IBV S species in cells and at the cell surface

The IBV S protein is cleaved by a furin-like protease generating the S1 and S2 subunits during trafficking through the Golgi, and at a second site (S2’) that primes the protein for fusion with the host cell (30). The S protein of IBV-EG3 is subject to further proteolysis near the junction of the protein with the viral envelope, resulting in a C-terminal fragment we term the ‘stub’ (22). We predicted that neutralization of Golgi pH by IBV E during infection protects the IBV S protein from premature proteolysis at the normal acidic pH of the trans-Golgi. We also predicted that processing and trafficking of IBV S in the presence of a functional pH altering viroporin (i.e. IBV E or IAV M2) would be similar, while a defunct viroporin (IBV EG3) would produce aberrant cleavage and more IBV S at the surface of cells. To test these predictions, we first demonstrated that when transiently overexpressed in Vero cells, IAV M2 neutralized the trans-Golgi in our pHluorin/ flow cytometry assay in the absence of the M2 inhibitor amantadine, but not in the presence of 5 μM amantadine as expected (Figure 4, (5–8)). With proof of principal established for M2 pH alteration during transfection, cells were cotransfected with IBV S and IBV E, or IBV EG3, with or without the IAV M2 protein. Surface biotinylation was performed, and the quantity of cleaved and uncleaved IBV S species in cell lysates and at the cell surface after streptavidin pull-down were determined by western blot analysis (Figure 5A). As predicted there was a significant increase in the total amount of IBV S at the surface of EG3 expressing cells as compared to WT IBV E. Notably, the presence of M2 in EG3 expressing cells reduced the amount IBV S at the surface of cells to a level comparable to cells expressing WT IBV E (Figure 5B.). Additionally, the total amounts of cleaved IBV S species (S2, S2’, and stub) were significantly increased in cell lysates expressing EG3 when compared to WT IBV E, and the presence of M2 in EG3 expressing cells reduced the amount of cleaved IBV S species back to levels comparable with cells expressing WT IBV E (Figure 5C). The same trend toward increased IBV S cleavage in EG3 expressing cells and a reduction in the cleaved species in the presence of M2 were also observed at the cell surface, though the differences were not significant.

**Figure 4.**
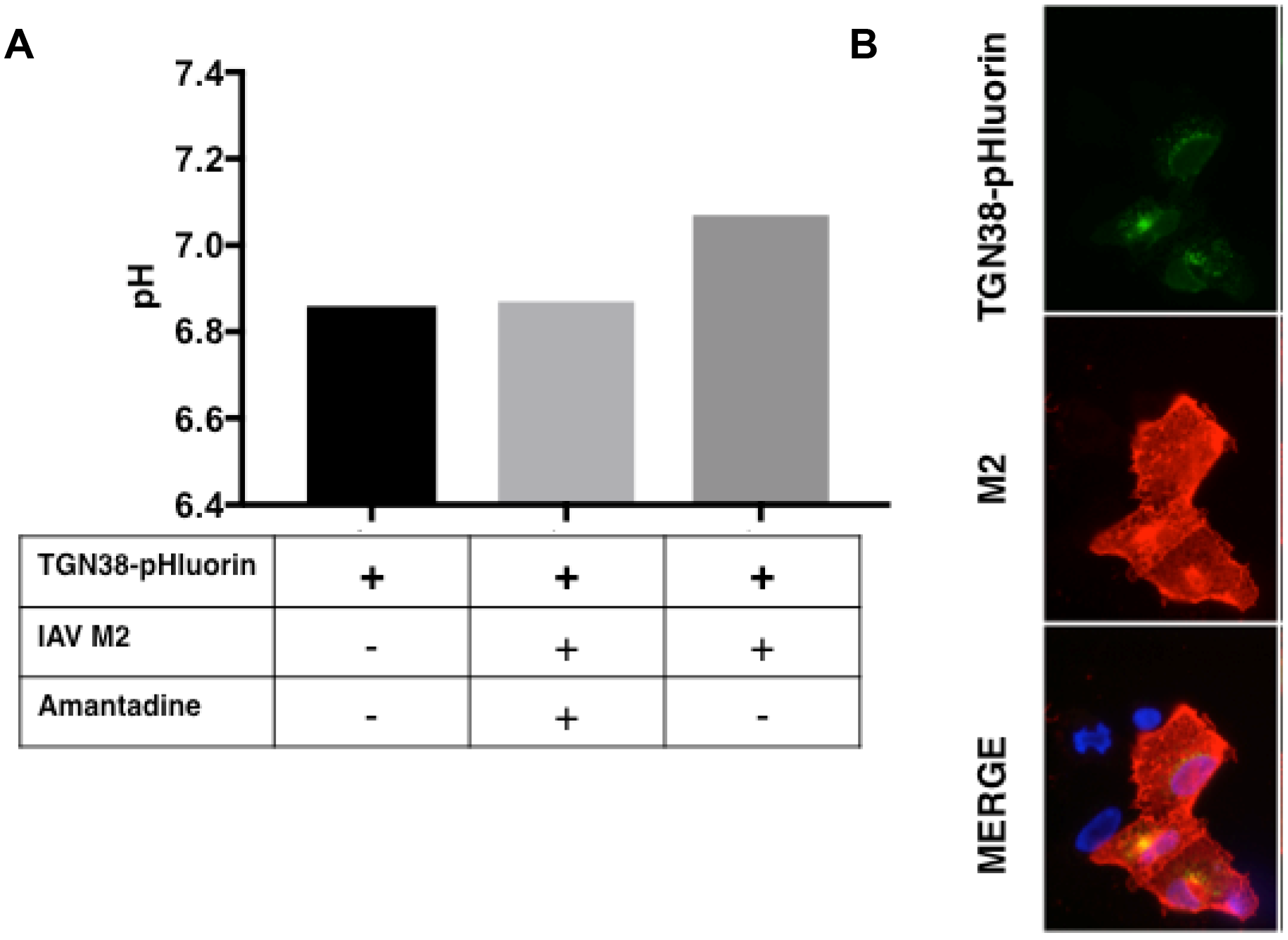
Influenza A M2 alters Golgi pH.(A) Vero cells transiently expressing the TGN38-pHluorin with or without transient expression of influenza M2 and with or without treatment with amantadine (5 μM) were evaluated by flow cytometry. The calculated pH values from a single independent experiment are graphed (~5,000 cells each).(B) Vero cells expressing the TGN38-pHluorin and M2 were evaluated by indirect immunofluorescence microscopy. Cells were labeled with rabbit anti-GFP and mouse anti-M2, followed by Alexa 488 anti-rabbit IgG and Alexa 546 anti-mouse IgG, and Hoescht stain.

**Figure 5.**
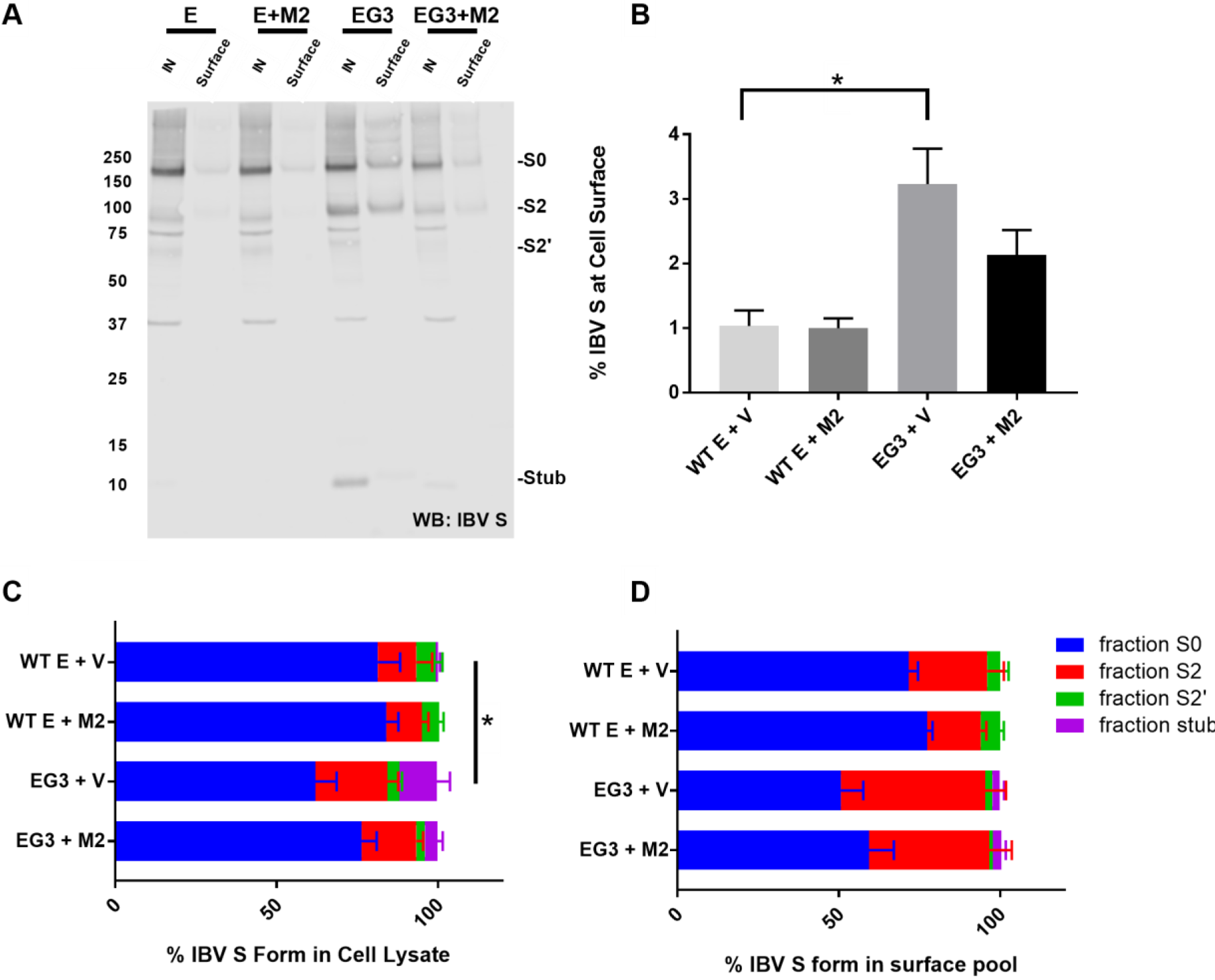
IAV M2 decreases the total amount cleaved IBV S species in cells and at the cell surface. (A) Vero cells expressing IBV S with either WT IBV E or EG3, along with either empty vector or IAV M2 were surface biotinylated‥Cells were lysed, and biotinylated proteins were isolated with streptavidin-agarose beads. Both input (10%) and surface fractions (100%) were subjected to western blot analysis with rabbit anti-IBV S followed by donkey anti-rabbit IgG-680 (LI-COR). IN = input. The positions of the IBV S species are indicated, as are the molecular weight markers. (B) The percent of total IBV S at the surface of cells was evaluated. The low percent of surface S is likely due to inefficient biotinylation. One-way ANOVA was performed with GaphPad Prism, * = p value <<0.05 when compared to WT E + vector. The fraction of each IBV S species in the lysates (C) and surface pool (D) was expressed as the percent of the total S. A Two-way ANOVA was performed with GraphPad Prism to evaluate the total amount of IBV S cleavage products in the input and surface fractions. *= p value <0.05 when compared to WT E + vector. Error bars = SEM.

## Discussion

### Neutralization of the Golgi by IBV E

Our flow cytometry and pH sensitive ratiometric analysis demonstrated that the Golgi luminal pH is increased in IBV infected cells. Overexpression of IBV E caused a similar increase in the pH of the lumen of the trans-Golgi and also increased the pH of the *medial*-Golgi. The average baseline measurement for TGN38-pHluorin in three independent experiments in which ~5000 cells were measured was pH 6.78. While the presence of overexpressed IBV M did not significantly increase the pH of the Golgi, the presence of overexpressed IBV E significantly increased the pH to 7.06, an increase of 0.28 pH units. In infected Vero cells where ~8,000 cells were analyzed per experiment, a pH increase of 0. 35 pH units from the uninfected baseline measurement of pH 6.76 to infected pH of 7.11 suggested the pH increase may be physiologically important. Our baseline measurement of pH 6.78 in the context of transient overexpression of the TGN38-pHluorin and IBV E protein in HeLa cells, and the baseline of 6.76 in the context of Vero cells stably expressing the TGN38-pHluorin are both similar, though somewhat higher, than pH values reported in the literature for the TGN. Values for the TGN38-pHluorin in different cell types range from pH 6.2-6.7 (28, 31, 32). We believe that the slightly higher baseline measurement we observed is likely due to some plasma membrane cycling of the TGN38-pHluorin, which would increase the average pH in a given cell since the pH of the extra cellular buffer was pH 7.3. The cycling of TGN38-pHlorin is likely to be the predominant reason for a higher than expected TGN pH, since the *medial*-Golgi pHluorin reported a pH of ~6.4. We believe that the consistent shift in the Golgi pH in cells expressing IBV E is more important than the actual baseline pH value we observed with the TGN38-pHlorin. Previously, the TGN38-pHluorin construct was used to demonstrate that CHO cells lacking the counter-ion channel Golgi pH regulator (GPHR) had a 0.4 pH unit increase, which substantiates the magnitude of the observed pH increase in this study.

### Neutralization of Golgi luminal pH correlates with secretory pathway disruption and the LMW pool of IBV E

Our analysis of the HD mutants of IBV E demonstrated that alkalization of the TGN induced by overexpression of wild-type IBV E correlates with the presence of the LMW, likely monomeric, pool of IBV E, which is the predominant form of the protein observed in cells expressing the IBV E^A26F^ mutant. The IBV E^A26F^ mutant elicited a pH shift of 0.43 units, slightly higher than the increase observed with the WT IBV E protein (0.28 pH unit increase). However, the IBV E^T16A^ or IBV EG3 HD mutants did not cause a statistically significant shift (0.20 and 0.12 pH units, respectively). We previously showed that the LMW pool of IBV E correlates with the secretory pathway disruption, including Golgi disassembly and slow trafficking of model cargo proteins, observed when the wild-type IBV E protein is expressed (23, 29). We hypothesized that the Golgi disruption was likely occurring in an IBV E ion channel-independent manner (26, 29). This study corroborates our hypothesis and suggests that the monomeric form of IBV E causes Golgi disruption via alteration of Golgi luminal pH through a mechanism involving interaction with a host protein. This interpretation is strengthened by evidence suggesting that neither the IBV E^T16A^ or IBV E^A26F^ mutants have ion channel activity in artificial membranes (27), suggesting that any remaining HMW weight, oligomeric IBV E that may be present when IBV E^A26F^ is expressed is not likely eliciting secretory pathway disruption or pH neutralization via IBV E ion channel activity. However, To *et al.* suggest that mutation of IBV E^T16A^ and not IBV E^A26F^ prevents oligomerization of IBV E, the opposite of our findings. The difference in these results is likely due to different modes of protein expression and the downstream assays used to evaluate oligomerization. In our studies we expressed the HD mutants in mammalian cells and evaluated oligomerization via sucrose gradient analysis followed by cross-linking and immunoprecipitation of gradient fractions of interest (26). To *et al.* bacterially expressed these proteins and then analyzed them via native PAGE electrophoresis after purifying and resolubilizing the samples (27).

### Neutralization of Golgi luminal pH by IBV E and IAV M2 correlates with reduced cleavage of IBV S protein

A proposed model for the cleavage and function of CoV S proteins suggests that proteolytic processing at two cleavage sites (S1/S2 and S2’) releases the protein from its prefusion conformation and allows for a conformational change that exposes the fusion peptide (33). This change in conformation may also release the S1 subunits from The S2 subunits of the CoV S trimer (33). A possible detrimental consequence of the normal acidic Golgi pH to the virus could be that the S protein is subject to a conformational change and premature and possibly excessive proteolytic processing, resulting in release of the S1 subunit prior to receptor binding. This would result in noninfectious or impaired virions. Here we demonstrate that when IBV S is overexpressed in cells in the presence of EG3, the levels of IBV S cleavage species are increased significantly as compared to IBV S in the presence of IBV E or IAV M2, lending support to the hypothesis that the neutralization of Golgi pH by IBV E protects IBV S from premature cleavage. Additionally, when over expressed in cells IBV S is present to a greater extent at the cell surface, corroborating previous data observed during IBV EG3 infection (23).

### CoV E protein viroporins and protein interaction

What is striking about IBV E when compared to M2 or p7 is the notion that IBV E is likely causing secretory pathway disruption in an IBV E ion-channel independent way. If IBV E is acting as a monomer to elicit secretory pathway disruption and alter pH, interaction of IBV E with a host protein(s) or lipid(s) would be necessary.

Evidence suggests that CoV E proteins interact with host cell proteins to influence pathogenicity. Interaction of SARS-CoV E with the PDZ (post-synaptic density protein-95/discs Large/zonula occludens-1) domain containing scaffolding protein, syntenin, and with the tight junction protein PALS1 (protein associated with Lin Seven 1) through its PBM (PDZ- binding motif) domain, implicates the E protein as a pathogenic determinant (34–39). The importance of the SARS-CoV E PBM domain was made more evident in studies addressing SARS-CoV infection with viruses lacking the E protein (40). During passage in tissue culture viruses lacking the E gene incorporated a partial duplication of the M gene in which novel and variable PBM domains were incorporated into the chimeric M gene (40). Similarly, deletion of the E gene in recombinant MHV has also been shown to result in partial duplication of M gene (41). When SARS-CoV lacking the E protein was passaged in mice a partial duplication of the 8a protein occurred, which contained a novel PBM domain in the chimeric 8a protein. It is worth noting that not all CoV E proteins have a PBM. In fact, IBV E lacks a PBM. It is tempting to speculate that relatively long C-terminus of IBV E compared to SARS-CoV E provides for the potential for host protein interaction in absence of a PBM domain. The differences in the length and relevant domains in the C-terminus of CoV E proteins are of interest in the field.

Interestingly, SARS-CoV 3a and 8a proteins have been shown to have ion-channel activity in artificial membranes in addition to E (39, 42–44). Intriguingly, the SARS-CoV 3a protein elicits inflammatory signaling similar to the SARS-CoV E protein, suggesting that CoVs may encode accessory proteins that i) may overlap in function to the E protein or ii) that may substitute for a particular role of the multifunctional E protein. Additionally, the SARS-CoV 3a protein has been show to induce Golgi fragmentation as an antagonist to the Arf1 GTPase involved in maintaining the structure and function of the Golgi (45). Further study of these SARS-CoV viroporins outside the complicated context of infection, especially their ability to induce pH changes in the lumen of the secretory pathway, may inform our studies on the IBV E protein.

Whether the endogenous role of putative IBV E interacting protein(s) be pH maintenance, vesicle formation, membrane architecture, protein glycosylation or any number of jobs performed by constituents of the secretory pathway, interaction of the protein(s) or lipid(s) with the large amount of IBV E known to reside at the ERGIC could disrupt the pH directly or indirectly by interfering with normal secretion or architecture of the Golgi. In summary, studies addressing CoV E protein-protein and protein-lipid interactions, and the development of tools to evaluate ion-channel activity *in vivo*, will go a long way in elucidating the precise mechanisms of the multifunctional family of viroporin proteins.

## Acknowledgements

We thank Helene Verhije (Department of Pathology, Utrecht University) for plasmids used in the construction of the codon-optimized IBV S construct and Andrew Pekosz (Department of Molecular Microbiology and Immunology, The Johns Hopkins Bloomberg School of Public Health) for M2 expression construct. We also thank the past and present members of the Machamer lab for critical and helpful discussions about the data presented here. This work was supported by NIH R01 GM117399 and NIH T32 GM007445.

## Materials and Methods

### Cell Culture

HeLa and Vero cells were cultured in Dulbecco’s modified Eagle medium (DMEM; Invitrogen/Gibco, Grand Island, NY) containing 10% fetal bovine serum (FBS; Atlanta Biologicals, Lawrenceville, GA) and 0.1 mg/ml Normocin (Invivogen, San Diego, CA) at 37°C under 5% CO_2_.

### Plasmids

The pCAGSS IBV E, IBV EG3, IBV M, IBV E^T16A^, IBV E^A26F^, codon optimized IBV S, M2 and empty pCAGGS-MCS plasmids have been previously described (23, 29, 46–49). The pCAGGS M2 plasmid was a generous gift from Dr. Andrew Pekosz. The pME-zeo-pHluorin-TGN38 plasmid and GnT1-pHluorin plasmids have been previously described and were generous gifts from Dr. Yusuke Maeda (28).

### Transient transfection

X-tremeGENE 9 DNA Transfection Reagent (Roche, Indianapolis, IN) was used to transiently transfect cells according to the manufacturer’s protocol. Subconfluent HeLa cells in 6 mm dishes were transfected with 1 μg of each plasmid indicated for a particular experiment, diluted into Opti-MEM (Invitrogen/Gibco) with a 1:3 ratio of X-tremeGENE 9. When the TGN38-pHluorin plasmid was transfected alone the pCAGGS-MCS empty vector was transfected to control for the tota
l amount of DNA transfected. For the cotransfection of IBV S with E, EG3 and IAV M2, subconfluent Vero cells in 6 well dishes were transfected with a total of 2 ug of DNA: 1 ug of pCAGGS/IBV S plus 0.5 ug pCAGGS/IBV E or EG3, and 0.5 ug pCAGGS/MCS or pCAGGS/IAV M2 as described above.

### Antibodies

The rabbit polyclonal antibodies recognizing the C-terminus of IBV E and the N-terminal head of golgin-160 have been previously described (50, 51). The antibody generated to the C-terminus of IBV S has also been described (52). The mouse monoclonal antibody recognizing the N-terminus of influenza A M2 has been previously described and was a generous gift from Dr. Andrew Pekosz (53). The mouse anti-GFP antibody was from Roche (Mannheim, Germany). The rabbit anti-GFP antibody was from Thermo Fisher Scientific (Rockford, IL). Alexa-Fluor 488-conjugated anti-rabbit IgG, Alexa-Fluor 488-conjugated anti-mouse IgG, Alexa Fluor 568-conjugated anti-rabbit IgG, and Alexa Fluor 568-conjugated anti-mouse IgG were from Invitrogen/Molecular Probes (Eugene, OR).

### Indirect immunofluorescence microscopy

Cells were washed with PBS and fixed in 3% paraformaldehyde in PBS for 10 min at 22°C. The fixative was quenched in PBS containing 10 mM glycine (PBS-Gly), and the cells were permeabilized in 0.5% Triton X-100 for 3 min. The coverslips were washed twice with PBS-Gly and incubated in primary antibody with 1% BSA for 20 min at room temperature. Rabbit anti-IBV E and rabbit anti-golgin160 were used at 1:1,000. Rabbit anti-GFP and mouse anti-M2 were used at 1:500. Mouse anti-GFP was used at 1:300. The cells were then washed twice with PBS-Gly and incubated for 20 min in secondary antibody with 1% BSA. Alexa-Fluor 488-conjugated anti-rabbit IgG and Alexa Fluor 568-conjugated antimouse IgG were used at 1:1,000. The coverslips were washed twice in PBS-Gly and incubated with Hoescht 33285 [0.1 μg/ml] to stain DNA, rinsed twice in PBS-Gly and mounted on slides in glycerol containing 0.1M *N*-propylgallate. Images were captured using an Axioskop microscope (Zeiss) equipped for epifluorescence with an ORCA-03G charge-coupled-device camera (Hamamatsu, Japan) and iVision software (Bio Vision Technologies).

### Establishment of TGN38-pHluorin stable Vero cell line

Subconfluent Vero cells were transfected with the pME-zeo-pHluorin-TGN38 plasmid according to the manufacturer’s protocol. Transfected cells were grown in DMEM containing 10% FBS under selection with Zeocin (Invitrogen) at 250 μg/ml. Individual clones were selected and evaluated by indirect immunofluorescence microscopy.

### Determination of Golgi pH during IBV infection

The Beaudette strain of recombinant IBV used in this study has been previously described (54). Vero cells stably expressing TGN38-pHluorin, seeded on 6 cm dishes at 3.5×10^5^, were inoculated with IBV diluted to an MOI of 25 in SF DMEM, and virus was adsorbed for 1 h with rocking. Inoculum was removed, and the cells were rinsed with DMEM containing 5% FBS. The cells were then incubated at 37°C in DMEM containing 5% FBS for 18 h. Cells were then washed with phosphate buffered saline (PBS) and trypsinized for 3 min and resuspended in ice cold SF DMEM. Cells were centrifuged at 112 × g and washed with ice cold SF DMEM twice. Cells were centrifuged as above and resuspended in 1 ml of the calibration buffers (140 mM KCl, 2 mM CaC_2_, 1mM MgSO_4_, 1.5 mM K_2_HPO_4_, 10 mM glucose, 10 mM MES, 10 mM HEPES, 10 μM monensin, 10 μM nigericin) of specified pH to generate a calibration curve, or Na-RINGER Buffer pH 7.3 (140 mM NaCl, 2 mM CaCh, 1mM MgSO_4_, 1.5 mM K_2_HPO_4_, 10 mM glucose, 10 mM MES, 10 mM HEPES) in the case of experimental samples. The cells were incubated at room temperature for ~10 min before being run through a Becton Dickinson LSRII flow cytometer. The pHluorin was excited at 405 nm and 488 nm and the emission signals were collected with detection filters at 500-550 nm and 515-545 nm, respectively. Flow cytometric data was collected and quantified using FACS Diva software 8.0. The emission ratios (405:488) of TGN38-pHluorin in calibration buffers of known pH were used to generate a linear calibration curve (Microsoft Excel) with which to calculate the luminal Golgi pH in infected and uninfected cells.

### Transient expression of TGN38-pHluorin or GnT1-pHluorin and IBV E or HD mutants in HeLa cells

At 12 h post-transfection, HeLa cells transiently expressing wild-type (WT) or mutant IBV E and TGN38-pHluorin or GnT1-pHluorin, or TGN38-pHluorin alone or GNT1-pHluorin alone, were washed with serum-free (SF) DMEM and incubated for 1 h at 37°C in SF DMEM containing 100 μg/ml cycloheximide. Cells were then washed with phosphate buffered saline (PBS) and trypsinized for 3 min and resuspended in ice cold SF DMEM. Cells were centrifuged at 112 × g and washed with ice cold SF DMEM twice. Cells were centrifuged as above and resuspended in 1 ml of the calibration buffers to generate the standard curve as described above, as well as in buffer lacking the ionophores to determine the Golgi pH. IBV M, another viral membrane protein localized to the Golgi, was used as a membrane protein overexpression control.

### Surface biotinylation and assessment of IBV S proteolytic processing

At 24h post-transfection, Vero cells expressing IBV S with either WT IBV E or EG3, along with either empty vector or IAV M2 were incubated with 0.5 mg/ml EZlink-NHS-SS-biotin (Pierce) in Hank’s buffered salt solution at 0°C for 30 min. After quenching the biotin in PBS with 10 mM glycine for 5 min at 0°C, cells were rinsed in PBS, scraped into PBS and pelleted at 4K RPM for 2.5 min. After lysis in 60 ul 1% NP40, 0.4% deoxycholate, 50 mM Tris-HCl pH 8.0, and 62.5 mM EDTA at 0°C for 20 min, samples were clarified by spinning at 14K RPM for 10 min at 4°C. Ten percent (6 ul) was removed for “input” (and combined with 6 ul of 2X sample buffer) and the remainder was diluted with 200 ul NHN (1% NP40, 10 mM Hepes pH 7.2, 150 mM NaCl). Streptavidin-agarose beads (50 ul of a 50% slurry, Pierce) were added and samples were incubated for 2h at 4°C and 30 min at 22°C. The beads were washed twice in NHN and eluted in 2x sample buffer with 5% p-mercaptoethanol at 85°C for 5 min. Samples were electrophoresed in NUPAGE 4-12% gradient gels (ThermoFisher) and transferred to low fluorescence PVDF (Millipore). After blocking in 10 mM Tris-HCl pH 7,4, 0.15 M NaCl (TBS) plus 5% nonfat milk, membranes were probed with rabbit anti-IBV S diluted 1:5000 in TBS/0.05% Tween (TBS/Tw). Bands were detected with donkey anti-rabbit IgG-680 (LI-COR) diluted in TBS/Tw, and imaged using the LI-COR Odyssey CLx. Quantification was performed in Image Studio (LI-COR), and each IBV S fragment was expressed as the percent of the total S for both the input and surface fractions.

